# Bayesian Inference of Natural Selection from Allele Frequency Time Series

**DOI:** 10.1101/037200

**Authors:** Joshua G. Schraiber, Steven N. Evans, Montgomery Slatkin

## Abstract

The advent of accessible ancient DNA technology now allows the direct ascertainment of allele frequencies in ancestral populations, thereby enabling the use of allele frequency time series to detect and estimate natural selection. Such direct observations of allele frequency dynamics are expected to be more powerful than inferences made using patterns of linked neutral variation obtained from modern individuals. We developed a Bayesian method to make use of allele frequency time series data and infer the parameters of general diploid selection, along with allele age, in non-equilibrium populations. We introduce a novel path augmentation approach, in which we use Markov chain Monte Carlo to integrate over the space of allele frequency trajectories consistent with the observed data. Using simulations, we show that this approach has good power to estimate selection coefficients and allele age. Moreover, when applying our approach to data on horse coat color, we find that ignoring a relevant demographic history can significantly bias the results of inference. Our approach is made available in a C++ software package.

## 1. INTRODUCTION

The ability to obtain high-quality genetic data from ancient samples is revolutionizing the way that we understand the evolutionary history of populations. One of the most powerful applications of ancient DNA (aDNA) is to study the action of natural selection. While methods making use of only modern DNA sequences have successfully identified loci evolving subject to natural selection [Nielsen et al., 2005, Voight et al., 2006, Pickrell et al., 2009], they are inherently limited because they look indirectly for selection, finding its signature in nearby neutral variation. In contrast, by sequencing ancient individuals, it is possible to directly track the change in allele frequency that is characteristic of the action of natural selection. This approach has been exploited recently using whole genome data to identify candidate loci under selection in European humans [Mathieson et al., 2015].

To infer the action of natural selection rigorously, several methods have been developed to explicitly fit a population genetic model to a time series of allele frequencies obtained via aDNA. Initially, Bollback et al. [2008] extended an approach devised by Williamson and Slatkin [1999] to estimate the population-scaled selection coefficient, *α* = 2*N_e_s*, along with the effective size, *N_e_*. To incorporate natural selection, Bollback et al. [2008] used the continuous diffusion approximation to the discrete Wright-Fisher model. This required them to use numerical techniques to solve the partial differential equation (PDE) associated with transition densities of the diffusion approximation to calculate the probabilities of the population allele frequencies at each time point. Ludwig et al. [2009] obtained an aDNA time series from 6 coat-color-related loci in horses and applied the method of Bollback et al. [2008] to find that 2 of them, ASIP and MC1R, showed evidence of strong positive selection.

Recently, a number of methods have been proposed to extend the generality of the Bollback et al. [2008] framework. To define the hidden Markov model they use, Bollback et al. [2008] were required to posit a prior distribution on the allele frequency at the first time point. They chose to use a uniform prior on the initial frequency; however, in truth the initial allele frequency is dictated by the fact that the allele at some point arose as a new mutation. Using this information, Malaspinas et al. [2012] developed a method that also infers allele age. They also extended the selection model of Bollback et al. [2008] to include fully recessive fitness effects. A more general selective model was implemented by Steinrücken et al. [2014], who model general diploid selection, and hence they are able to fit data where selection acts in an over- or under-dominant fashion; however, Steinrücken et al. [2014] assumed a model with recurrent mutation and hence could not estimate allele age. The work of Mathieson and McVean [2013] is designed for inference of metapopulations over short time scales and so it is computationally feasible for them to use a discrete time, finite population Wright-Fisher model. Finally, the approach of Feder et al. [2014] is ideally suited to experimental evolution studies because they work in a strong selection, weak drift limit that is common in evolving microbial populations.

One key way that these methods differ from each other is in how they compute the probability of the underlying allele frequency changes. For instance, Malaspinas et al. [2012] approximated the diffusion with a birth-death type Markov chain, while Steinrücken et al. [2014] approximate the likelihood analytically using a spectral representation of the diffusion discovered by Song and Steinrücken [2012]. These different computational strategies are necessary because of the inherent difficulty in solving the Wright-Fisher partial differential equation. A different approach, used by Mathieson and McVean [2013] in the context of a densely-sampled discrete Wright-Fisher model, is to instead compute the probability of the entire allele frequency trajectory in between sampling times.

In this work, we develop a novel approach for inference of general diploid selection and allele age from allele frequency time series obtained from aDNA. The key innovation of our approach is that we impute the allele frequency trajectory between sampled points when they are sparsely-sampled. Moreover, by working with a diffusion approximation, we are able to easily incorporate general diploid selection and changing population size. This approach to inferring parameters from a sparsely-sampled diffusion is known as high-frequency path augmentation, and has been successfully applied in a number of contexts [Roberts and Stramer, 2001, Golightly and Wilkinson, 2005, 2008, Sprensen, 2009, Fuchs, 2013]. The diffusion approximation to the Wright-Fisher model, however, has several features that are atypical in the context of high-frequency path augmentation, including a time-dependent diffusion coefficient and a bounded state-space. We test this approach with simulation, showing that it’s important to accurately model demography history, then apply it to several datasets and find that we have power to estimate parameters of interest from real data.

## 2. MODEL AND METHODS

### 2.1 Overview

We begin by first reviewing the Wright-Fisher model, presenting its diffusion approximation as a stochastic differential equation (SDE). We then describe our inferential strategy using a path augmentation approach, in which we model the underlying allele frequency trajectory as an additional (infinite dimensional) parameter. This requires us to derive an expression for the likelihood of an allele frequency trajectory, including accounting for the fact that we model alleles that start from low frequency as new mutants. Finally, we describe a Markov chain Monte Carlo algorithm for obtaining a posterior distribution of the parameters of natural selection, as well as the allele frequency trajectory.

### 2.2 Generative model

We assume a randomly mating diploid population that is size *N(t)* at time *t*, where *t* is measured in units of 2*N*_0_ generations for some arbitrary, constant N_0_. At the locus of interest, the ancestral allele, *A*_0_, was fixed until some time t_0_ when the derived allele, *A*_1_, arose with diploid fitnesses as given in Table 1.

**TABLE 1.** Fitness scheme assumed in the text.

| Genotype | $A_1A_1$ | $A_1A_0$ | $A_0A_0$ |
| --- | --- | --- | --- |
| Fitness | $1 + s_2$ | $1 + s_1$ | 1 |

Given that an allele arises at some finite population frequency 0 < *x*_0_ < 1 at some time *t*_0_, the trajectory of population frequencies of *A*_1_ at times *t* ≥ *t*_0_, (*X_t_*)_*t*≥*t*_0__, is modeled by the usual diffusion approximation to the Wright-Fisher model (and many other models such as the Moran model), which we will henceforth call the Wright-Fisher diffusion. While many treatments of the Wright-Fisher diffusion define it in terms of the partial differential equation that characterizes its transition densities (e.g. Ewens [2004]), we instead describe it as the solution of a stochastic differential equation (SDE). Specifically, (*X_t_*)_*t*≥*t*_0__ satisfies the SDE

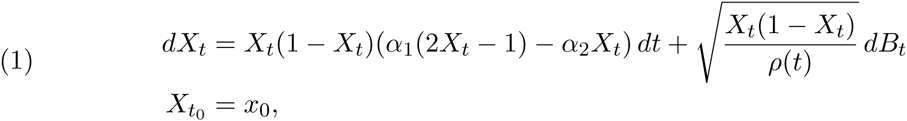

where *B* is a standard Brownian motion, *α*_1_ = 2*N*_0_*s*_1_, *α*_2_ = 2*N*_0_*s*_2_, and *ρ(t)* = *N(t)/N*_0_. If *X*_*t*_*__ = 0 (resp. *X*_*t*_*__ = 1) at some time *t*_*_ > *t*_0_, then *X_t_* = 0 (resp. *X_t_* = 1) for all *t* ≥ *t*_*_.

In order to make this description of the dynamics of the population allele frequency trajectory (*X_t_*)_*t*≥*t*_0__ complete, we need to specify an initial condition at time *t*_0_. In a finite population Wright-Fisher model we would take the allele *A*_1_ to have frequency 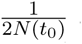 at the time t_0_ when it first arose in a single chromosome. This frequency converges to 0 when we pass to the diffusion limit, but we cannot start the Wright-Fisher diffusion at 0 at time *t*_0_ because the diffusion started at 0 remains at 0. Instead, we take the value of *X*_*t*0_ to be some small, but arbitrary, frequency *x*_0_. This arbitrariness in the choice of x_0_ may seem unsatisfactory, but we will see that, in the context of a Bayesian inference procedure, the resulting posterior distribution for the parameters *α*_1_, *α*_2_, *t*_0_ converges as *x*_0_ ↓ 0 to a limit which can be thought of as the posterior corresponding to a certain improper prior distribution, and so, in the end, there is actually no need to specify *x*_0_.

Finally, we require a model for how alleles arise. We assume that mutations at time *t* occur at a rate proportional to 2*N(t)*, and that a mutant allele arises exactly once. Further constraining alleles to have arisen more recently than some time, *T*, in the past, this implies that the prior density of allele ages is

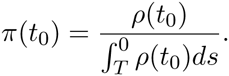

Taking the limit as *T* ↓ —∞ results in an improper distribution on allele age, which, in the context of our Bayesian inference algorithm, implies an improper prior distribution on *t*_0_ that is proportional to *ρ*. However, we emphasize that this still produces a proper posterior distribution on allele age (see also Slatkin [2001]).

Finally, we model the data assuming that at known times *t*_1_,*t*_2_,…, *t_k_* samples of known sizes *n*_1_, *n*_2_,…, *n_k_* chromosomes are taken and *c*_1_, *c*_2_,…, *c_k_* copies of the derived allele are found at the successive time points (Figure 1). Note that it is possible that some of the sampling times are more ancient than *t*_0_, the age of the allele.

**FIGURE 1.**
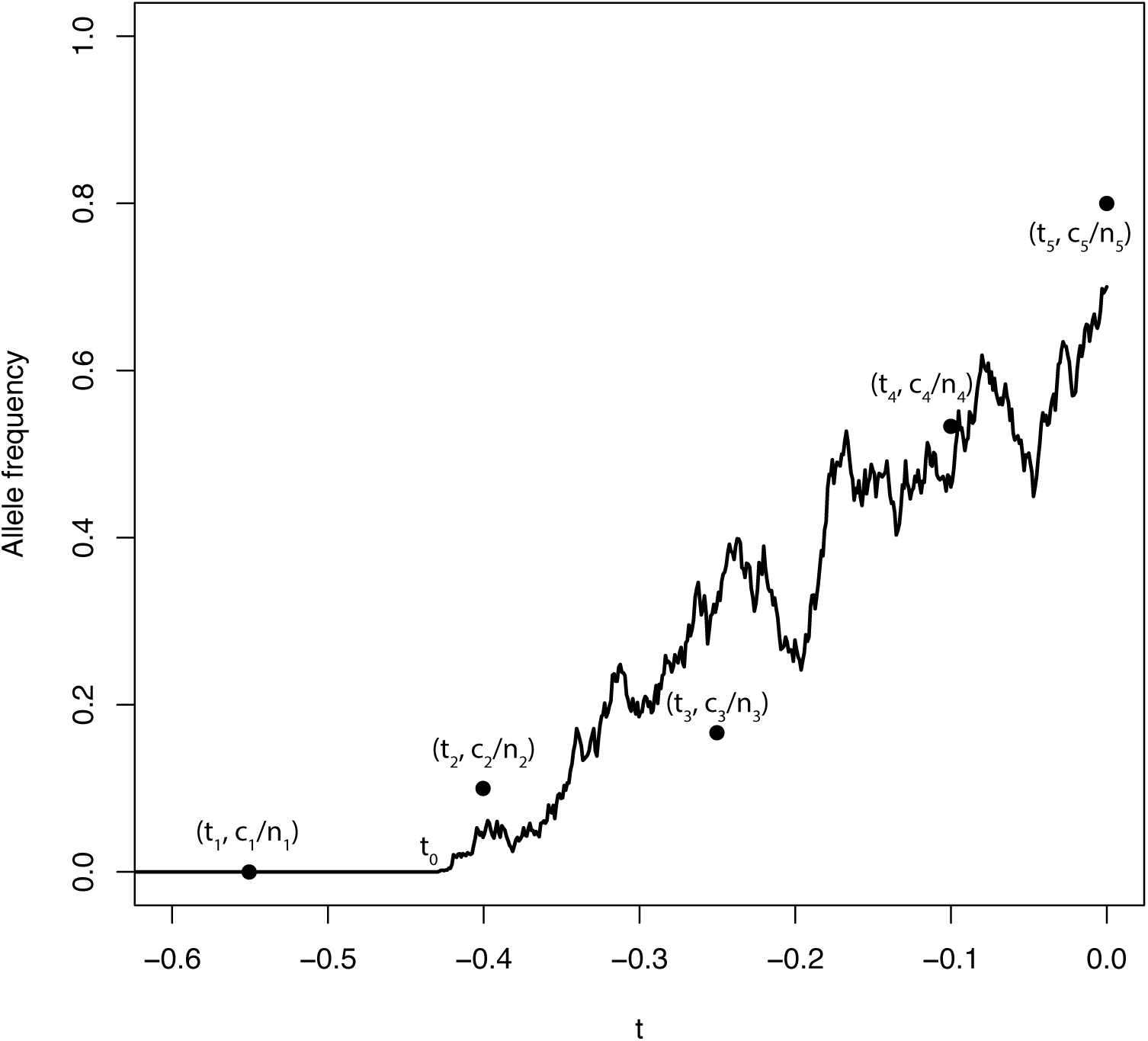
Taking samples from an allele frequency trajectory. An allele frequency trajectory is simulated from the Wright-Fisher diffusion (solid line). At each time, *t_i_* a sample of size *n_i_* chromosomes is taken and *c_i_* copies of the derived allele are observed. Each point corresponds to the observed allele frequency of sample *i.* Note that *t*_1_ is more ancient than the allele age, *t*_0_.

### 2.3 Bayesian path augmentation

We are interested in devising a Bayesian method to obtain the posterior distribution on the parameters, *α*_1_, *α*_2_, and *t*_0_ given the sampled allele frequencies and sample times – data which we denote collectively as D. Because we are dealing with objects that don’t necessarily have distributions which have densities with respect to canonical reference measures, it will be convenient in the beginning to treat priors and posteriors as probability measures rather than as density functions. For example, the posterior is the probability measure

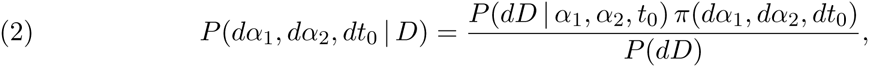

where *π* is a joint prior on the model parameters. However, computing the likelihood *P*(*dD* | *α*_1_, *α*_2_, *t*_0_) is computationally challenging because, implicitly,

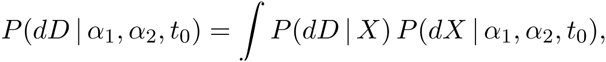

where the integral is over the (unobserved, infinite-dimensional) allele frequency path *X* = (*X_t_*)_*t*≥*t*_0__, *P*(· | *α*_1_, *α*_2_, *t*_0_) is the distribution of a Wright-Fisher diffusion with selection parameters *α*_1_, *α*_2_ started at time *t*_0_ at the small but arbitrary frequency *x*_0_, and

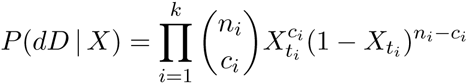

because we assume that sampled allele frequencies at the times *t*_1_,…, *t_k_* are independent binomial draws governed by underlying population allele frequencies at the these times. Integrating over the infinite-dimensional path (*X_t_*)_*t*≥*t*_0__ involves either solving partial differential equations numerically or using Monte Carlo methods to find the joint distribution of population allele frequency path at the times *t*_1_,…, *t_k_*.

To address this computational difficulty, we introduce a path augmentation method that treats the underlying allele frequency path (*X_t_*)_*t*≥*t*_0__ as an additional parameter. Observe that the posterior may be expanded out to

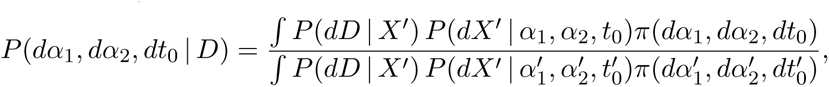

where we use primes to designate dummy variables over which we integrate. Thinking of the path (*X_t_*)_*t*≥*t*_0__ as another parameter and taking the prior distribution for the augmented family of parameters to be

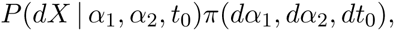

the posterior for the augmented family of parameters is

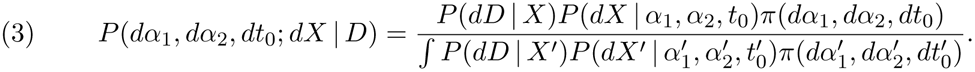

We thus see that treating the allele frequency path as a parameter is consistent with the initial “naive” Bayesian approach in that if we integrate the path variable out of the posterior (3) for the augmented family of parameters, then we recover the posterior (2) for the original family of parameters. In practice, this means that marginalizing out the path variable from a Monte Carlo approximation of the augmented posterior gives a Monte Carlo approximation of the original posterior.

Implicit in our set-up is the initial frequency *x*_0_ at time *t*_0_. Under the probability measure governing the Wright-Fisher diffusion, any process started from *x*_0_ = 0 will stay there forever. Thus, we would be forced to make an arbitrary choice of some *x*_0_ > 0 as the initial frequency of our allele. However, we argue in the Appendix that in the limit as *x*_0_ ↓ 0, we can achieve an improper prior distribution on the space of allele frequency trajectories. We stress that our inference using such an improper prior is not one that arises directly from a generative probability model for the allele frequency path. However, it does arise as a limit as the initial allele frequency *x*_0_ goes to zero of inferential procedures based on generative probability models and the limiting posterior distributions are probability distributions. Therefore, the parameters *α*_1_, *α*_2_, *t*_0_ retain their meaning, our conclusions can be thought of approximations to those that we would arrive at for all sufficiently small values of *x*_0_, and we are spared the necessity of making an arbitrary choice of *x*_0_.

### 2.4 Path likelihoods

Most instances of Bayesian inference in population genetics have hitherto involved finite-dimensional parameters. Recall that for continuous, finite-dimensional parameters, one simply includes the prior *density* of the parameter value in place of the prior *probability.* Finite dimensional parameters usually have densities defined with respect to Lebesgue measure in an appropriate dimension; however, there is no infinite-dimensional Lebesgue measure against which to define a density for our infinite-dimensional augmented path. We thus require a reference measure on the infinite-dimensional space of paths that will play a role analogous to that of Lebesgue measure in the finite-dimensional case, allowing us to write down the probability density for each sampled path.

To see what is involved, suppose we have a diffusion process (*Z_t_*)_*t*≥*t*_0__ that satisfies the SDE

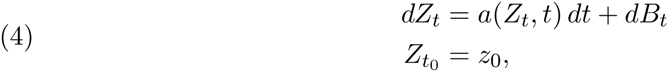

where *B* is a standard Brownian motion (the Wright-Fisher diffusion is not of this form but, as we shall soon see, it can be be reduced to it after suitable transformations of time and space). Let ℙ be the distribution of (*Z_t_*)_*t*≥*t*_0__ – this is a probability distribution on the space of continuous paths that start from position *z*_0_ at time *t*_0_. While the probability assigned by ℙ to any particular path is zero, we can, under appropriate conditions, make sense of the probability of a path under ℙ *relative* to its probability under the distribution of Brownian motion. If we denote by 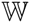 the distribution of Brownian motion starting from position *z*_0_ at time *t*_0_, then Girsanov’s theorem [Girsanov, 1960] gives the density of the path segment *(Z_s_*)_*t*_0_≤*s*≤*t*_ under ℙ relative to 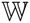 as

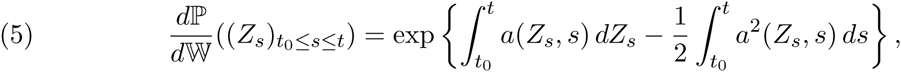

where the first integral in the exponentiand is an Itô integral. In order for (5) to hold, the integral 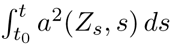 ds must be finite, in which case the Itô integral 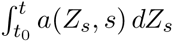 is also well-defined and finite.

However, the Wright-Fisher SDE (1) is not of the form (4). In particular, the factor multiplying the infinitesimal Brownian increment *dB_t_* (the so-called diffusion coefficient) depends on both space and time. To deal with this issue, we first apply a well-known time transformation (see e.g. Slatkin and Hudson [1991] and Griffiths and Tavare [1994]) and consider the process 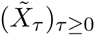 given by 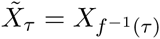, where

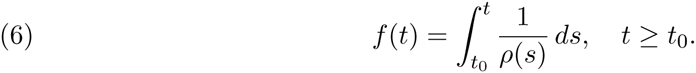

It is not hard to see that 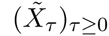 satisfies the following SDE with a time-independent diffusion coefficient,

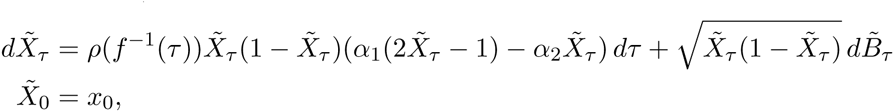

where 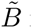 is a standard Brownian motion. Next, we employ an angular space transformation first suggested by Fisher [1922], 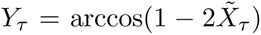. Applying Itô’s lemma [Itô, 1944] shows that (*Υ_τ_*)_*τ*≥0_ is a diffusion that satisfies the SDE

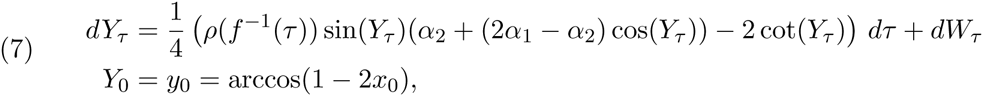

where *W* is a standard Brownian motion. If the process *X* hits either of the boundary points 0,1, then it stays there, and the same is true of the time and space transformed process *Y* for its boundary points 0, *π*.

The restriction of the distribution of the time and space transformed process *Y* to some set of paths that don’t hit the boundary is absolutely continuous with respect to the distribution of standard Brownian motion restricted to the same set; that is, the distribution of *Y* restricted to such a set of paths has a density with respect to the distribution of Brownian motion restricted to the same set. However, the infinitesimal mean in (7) (that is, the term multiplying *d_τ_*) becomes singular as *Y_τ_* approaches the boundary points 0 and *π*, corresponding to the boundary points 0 and 1 for allele frequencies. These singularities prevent the process *Y* from re-entering the interior of its state space and ensure that a Wright-Fisher path will be absorbed when the allele is either fixed or lost. A consequence is that the density of the distribution of *Y* relative to that of a Brownian motion blows up as the path approaches the boundary. We are modeling the appearance of a new mutation in terms of a Wright-Fisher diffusion starting at some small initial frequency *x*_0_ at time to and we want to perform our parameter inference in such a way that we get meaningful answers as *x*_0_ ↓ 0. This suggests that rather than working with the distribution 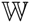 of Brownian motion as a reference measure it may be more appropriate to work with a tractable diffusion process that exhibits similar behavior near the boundary point 0.

To start making this idea of matching singularities more precise, consider a diffusion process 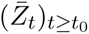 that satisfies the SDE

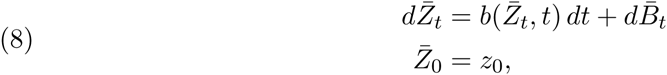

where 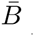 is a standard Brownian motion. Write ℚ for the distribution of the diffusion process 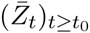 and recall that ℙ is the distribution of a solution of (4). If (*Z_s_*)_*t*_0_≤*s*≤*t*_ is a segment of path such that both 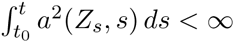 and 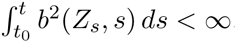 then

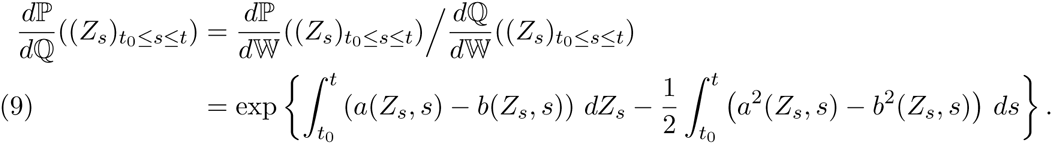

Note that the right-hand side will stay bounded if one considers a sequence of paths, indexed by *η* 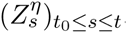, with 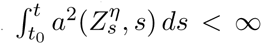 and 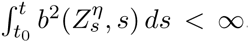, provided that 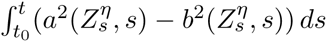 stays bounded. These manipulations with densities may seem somewhat heuristic, but they can be made rigorous and, moreover, the form of 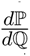 follows from an extension of Girsanov’s theorem that gives the density of ℙ with respect to ℚ directly without using the densities with respect to 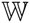 as intermediaries (see, for example, [Kallenberg, 2002, Theorem 18.10]).

We wish to apply this observation to the time and space transformed Wright-Fisher diffusion of (7). Because

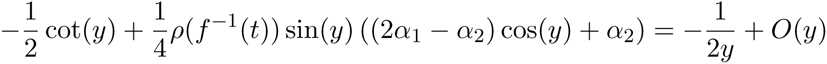

when *y* is small, an appropriate reference process should have infinitesimal mean *b(y, t)* ≈ —1/(2*y*) as *y* ↓ 0. Following suggestions by Schraiber et al. [2013] and Jenkins [2013], we compute path densities relative to the distribution ℚ of the Bessel(0) process, a process which is the solution of the SDE

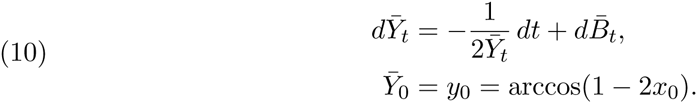

up until the first time that 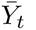 hits 0, after which time 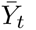 stays at 0 [Revuz and Yor, 1999, Chapter XI].

As we show more explicitly in the Appendix, this choice of dominating measure allows us to arrive at a proper posterior distribution as we send the initial frequency of the allele down to 0. In brief, if we write ℙ^*y*0^ and ℚ^*y*0^ for the respective distributions of the solutions of (7) and (10) to emphasize the dependence on *y*_0_ (equivalently, on the initial allele frequency *x*_0_), then there are *σ*-finite measures ℙ^0^ and ℚ^0^ with infinite total mass such that for each *ε* > 0

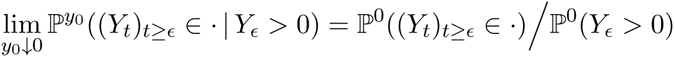

and

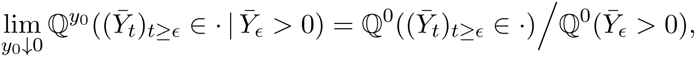

where the numerators and denominators in the last two equations are all finite. Moreover, ℙ^0^ has a density with respect to ℚ^0^ that arises by naively taking limits as *y*_0_ ↓ 0 in the functional form of the density of ℙ;^*y*0^ with respect to ℚ;^*y*0^ (we say “naively” because ℙ;^*y*0^ and ℚ;^*y*0^ assign all of their mass to paths that start at position *y*_0_ = arccos(1 − 2*x*_0_) at time 0, whereas ℚ^0^ and ℙ^0^ assign all of their mass to paths that start at position 0 at time 0, and so the set of paths at which it is relevant to compute the density changes as *y*_0_ ↓ 0). As we have already remarked, the limit of our Bayesian inferential procedure may be thought of as Bayesian inference with an improper prior, but we stress that the resulting posterior is proper.

The notion of the infinite measure ℚ^0^ may seem somewhat forbidding, but this measure is characterized by the following simple properties:

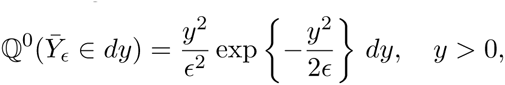

so that 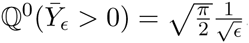, and conditional on the event 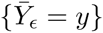 the evolution of 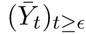 is exactly that of the Bessel(0) process started at position *y* at time *ε*. In the Appendix, we provide a more explicit construction of the measure ℚ^0^ as part of our derivation of the proposal ratios in our MCMC algorithm. Moreover, conditional on the event 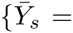 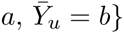 for 0 ≤ *s* < *u* and *a, b* > 0, the evolution of the “bridge” 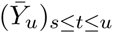 is the same as that of the corresponding bridge for a Bessel(4) process; a Bessel(4) process satisfies the SDE

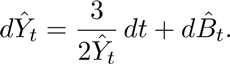

Very importantly for the sake of simulations, the Bessel(4) process is just the radial part of a 4-dimensional standard Brownian motion – in particular, this process started at 0 leaves immediately and never returns.

Note that the Bessel(0) process arises naturally because our space transformation *x* ↣ arccos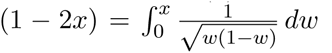 is approximately 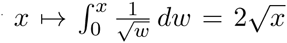 when *x* > 0 is small. Interestingly, a multiple of the square of Bessel(0) process, sometimes called Feller’s continuous state branching processes, arises naturally as an approximation to the Wright-Fisher diffusion for low frequencies and has a long history in population genetics [Haldane, 1927, Feller, 1951].

### 2.5 The joint likelihood of the data and the path

To write down down the full likelihood of the observations and the path, we make the assumption that the population size function *ρ(t)* is continuously differentiable except at a finite set of times *d*_1_ < *d*_2_ <… < *d_M_*. Further, we require that that 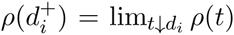 exists and is equal to *ρ(d_i_)* while 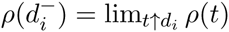 also exists (though it may not necessarily equal *ρ(d_i_)*).

We can write the joint likelihood of the data and the path as

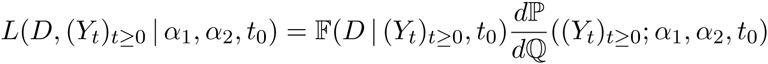

where 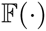 is the binomial sampling probability of the observed allele frequencies, ℙ is the distribution of transformed Wright-Fisher paths, and ℚ is the distribution of Bessel(0) paths. In the Appendix, we show that

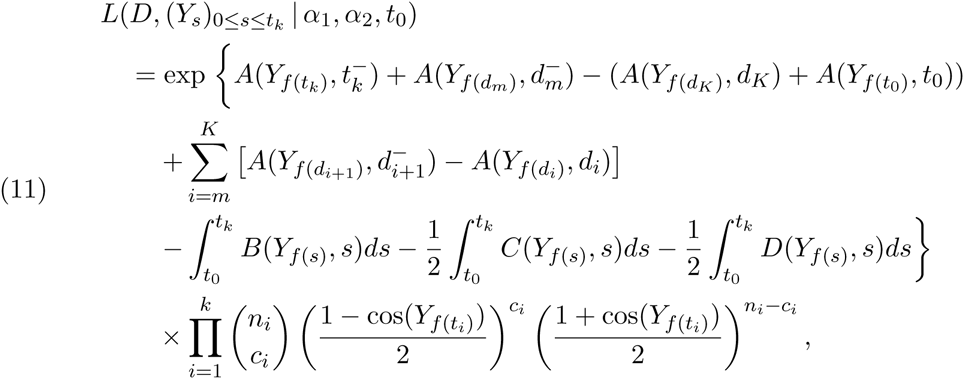

where *f* as in (6), *m* = min{*i*: *d_i_* > *t*_0_} and *K* = max{*i*: *d_i_* > *t_k_*}, and

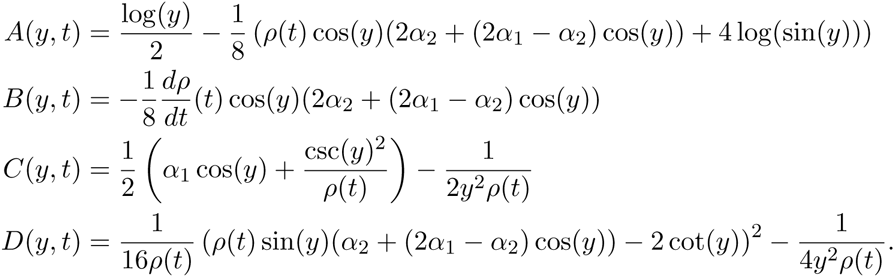

While this expression may appear complicated, it has the important feature that, unlike the form of the likelihood that would arise by simply applying Girsanov’s theorem, it only involves Lebesgue (indeed Riemann) integrals and not Itô integrals, which, as we recall in the Appendix, are known from the literature to be potentially difficult to compute numerically.

### 2.6 Metropolis-Hastings algorithm

We now describe a Markov chain Monte Carlo method for Bayesian inference of the parameters *α*_1_, *α*_2_ and *t*_0_, along with the allele frequency path (*X_t_*)_*t*≥*t*_0__ (equivalently, the transformed path (*Y_t_*)_*t*≥0_. While updates to the selection parameters *α*_1_ and *α*_2_ do not require updating the path, updating the time *t*_0_ at which the derived allele arose requires proposing updates to the segment of path from *t*_0_ up to the time of the first sample with a non-zero number of derived alleles. Additionally, we require proposals to update small sections of the path without updating any parameters and proposals to update the allele frequency at the most recent sample time.

#### 2.6.1. Interior path updates

To update a section of the allele frequency, we first choose a time *s*_1_ ∈ (*t*_0_,*t_k_*) uniformly at random, and then choose a time *s*_2_ that is a fixed fraction of the path length subsequent to *s*_1_. We prefer this approach of updating a fixed fraction of the path to an alternative strategy of holding *s*_2_ – *s*_1_ constant because paths for very strong selection may be quite short. Recalling the definition of f from (6), we subsequently propose a new segment of transformed path between the times *f* (*s*_1_) and *f* (*s*_2_) while keeping the values *Y_f_*(*s*_1_) and *Y_f_*(*s*_2_) fixed (Figure 2a). Such a path that is conditioned to take specified values at both end-points of the interval over which it is defined is called a bridge, and by updating small portions of the path instead of the whole path at once, we are able to obtain the desirable behavior that our Metropolis-Hastings algorithm is able to stay in regions of path space with high posterior probability. If we instead drew the whole path each time, we would much less efficiently target the posterior distribution.

**FIGURE 2.**
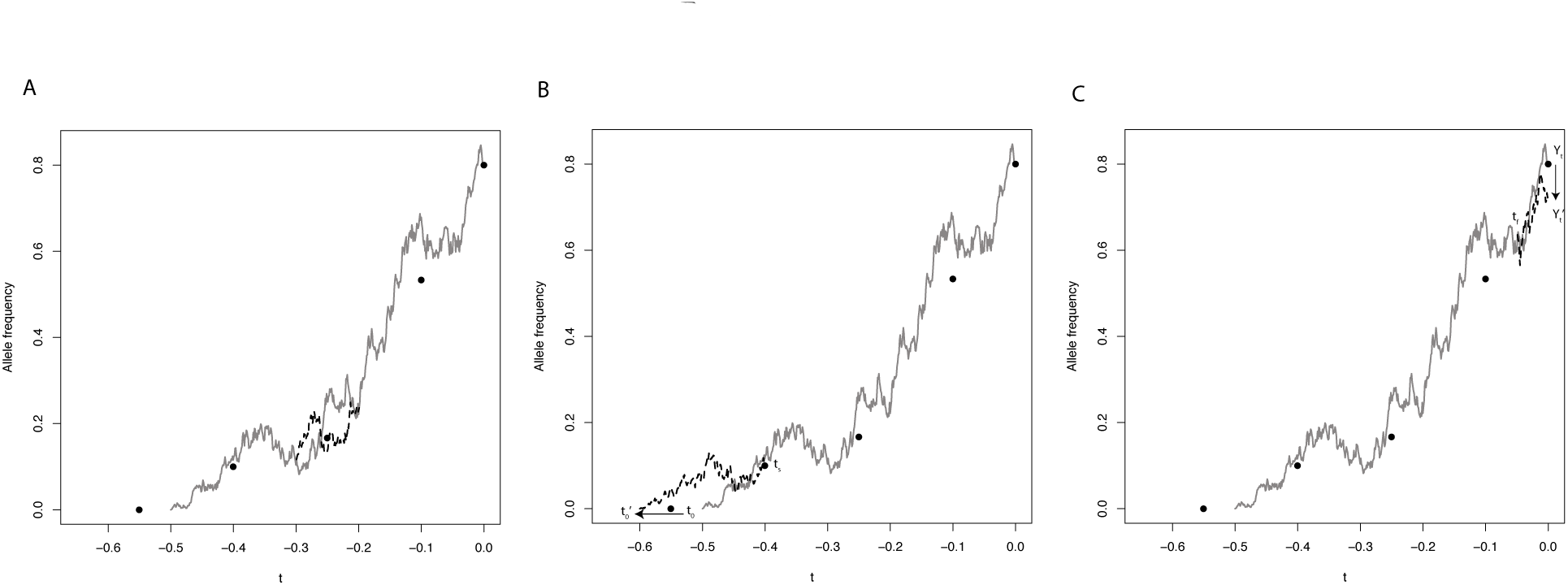
Illustration of path updates. Filled circles correspond to the same sample frequencies as in Figure 1. The solid gray line in each panel is the current allele frequency trajectory and the dashed black lines are the proposed updates. In panel a, an interior section of path is proposed between points *s*_1_ and *s*_2_. In panel b, a new allele age, 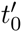 is proposed and a new path is drawn between 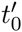 and *t_s_*. In panel c, a new most recent allele frequency 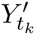 is proposed and a new path is drawn between *t_f_* and *t_k_*.

Noting that bridges must be sampled against the *transformed* time scale, the best bridges for the allele frequency path would be realizations of Wright-Fisher bridges themselves. However, sampling Wright-Fisher bridges is challenging (but see Schraiber et al. [2013], Jenkins and Spano [2015]), so we instead opt to sample bridges for the transformed path from the Bessel(0) process. Sampling Bessel(0) bridges can be accomplished by first sampling Bessel(4) bridges (as described in Schraiber et al. [2013]) and then recognizing that a Bessel(4) process is the same as a Bessel(0) process conditioned to never hit 0 and hence has the same bridges – in the language of the general theory of Markov processes, the Bessel(0) and Bessel(4) processes are Doob h-transforms of each other and it is well-known that processes related in this way share the same bridges. We denote by 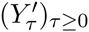 the path that has the proposed bridge spliced in between times *f* (*s*_1_) and *f* (*s*_2_) and coincides with (*Y_τ_*)_*τ*≥°_ outside the interval [*f*(*s*_1_), *f*(*s*_2_)].

In the Appendix, we show that the acceptance probability in this case is simply

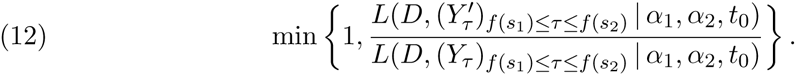

Note that we only need to compute the likelihood ratio for the segment of transformed path that changed between the times *f* (*s*_1_) and *f* (*s*_2_).

#### 2.6.2. Allele age updates

The first sample time with a non-zero count of the derived allele (Figure 2b) is *t_s_*, where *s* = min{*i: c_i_* > 0}. We must have *t*_0_ < *t_s_*. Along with proposing a new value 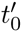 of the allele age *t*_0_ we will propose a new segment of the allele frequency path from time 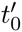 to time *t_s_*. Changing the allele age *t*_0_ to some new proposed value 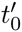 changes the definition of the function *f* in (6). Write 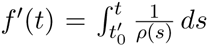, where we stress that the prime does not denote a derivative. The proposed transformed path *Y*’ consists of a new piece of path that goes from location 0 at time 0 to location *Y*_*f(t_s_)*_ at time *f*’(*t_s_*) and then has 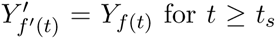. Recall that we use the improper prior *ρ(*t*_0_)* for *t*_0_, which reflects the fact that an allele is more likely to arise during times of large population size [Slatkin, 2001]. In the Appendix, we show that the acceptance probability is

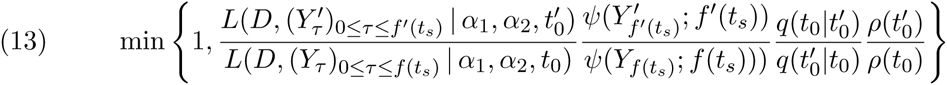

where, in the notation of Subsection 2.4,

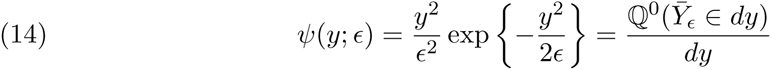

is the density of the so-called entrance law for the Bessel(0) process that appears in the characterization of the *σ*-finite measure ℚ^0^ and 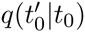 is the proposal distribution of 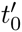 (in practice, we use a half-truncated normal distribution centered at *t*_0_, with the upper truncation occurring at the first time of non-zero observed allele frequency).

#### 2.6.3. Most recent allele frequency update

While the allele frequency at sample times *t*_1_,*t*_2_,…, *t*_*k*-1_ are updated implicitly by the interior path update, we update the allele frequency at the most recent sample time *t_k_* separately (note that the most recent allele frequency is not an additional parameter, but simply a random variable with a distribution implied by the Wright-Fisher model on paths). We do this by first proposing a new allele frequency 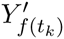 and then proposing a new bridge from *Y*_*f(t_f_)*_ to 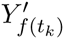 where *t_f_* ∈ (*t*_*k*−1_,*t_k_*) is a fixed time (Figure 2c). If 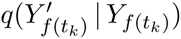 is the proposal density for 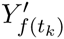 given *Y*_*f(t_k_)*_ (in practice, we use a truncated normal distribution centered at *Y*_*f(t_k_)*_ and truncated at 0 and π), then, arguing along the same lines as the interior path update and the allele age update, we accept this update with probability

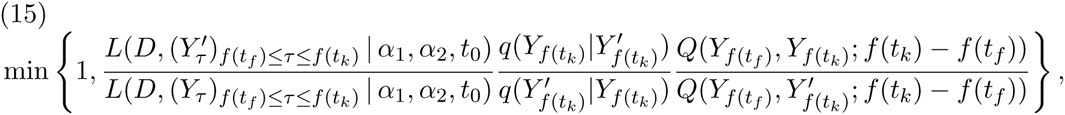

where

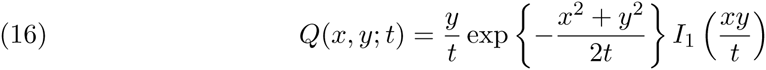

is the transition density of the Bessel(0) process (with *I*_1_(·) being the Bessel function of the first kind with index 1) – see Knight [1981, Section 4.3.6]. Again, it is only necessary to compute the likelihood ratio for the segment of transformed path that changed between the times *f*(*t_f_*) and *f*(*t_k_*).

### 2.7 Updates to *α*_1_ and *α*_2_

Updates to *α*_1_ and *α*_2_ are conventional scalar parameter updates. For example, letting 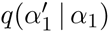 be the proposal density for the new value of *α*_1_, we accept the new proposal with probability

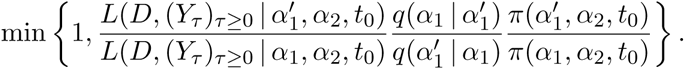

The acceptance probability for *α*_2_ is similar. For both *α*_1_ and *α*_2_, we use a heavy-tailed Cauchy prior with median 0 and scale parameter 100, and we take the parameters *α*_1_, *α*_2_, *t*_0_ to be independent under the prior distribution. In addition, we use a normal proposal distribution, centered around the current value of the parameter. Here, it is necessary to compute the likelihood across the whole path.

## 3. RESULTS

We first test our method using simulated data to assess its performance and then apply it to two real datasets from horses.

### 3.1 Simulation performance

To test the accuracy of our MCMC approach, we performed two sets of simulations. First, we simulated data under a constant demographic history to asses the quality of parameter inference under a simple model. Second, we simulated data under the horse demographic history of Der Sarkissian et al. [2015] and compared inferences performed with and without accounting for the demographic history.

In the constant demography simulations, we simulated allele frequency trajectories with ages uniformly distributed between 0.1 and 0.3 diffusion time units ago, evolving with *α*_1_ and *α*_2_ uniformly distributed between 0 and 100. We simulate allele frequency trajectories using an Euler approximation to the Wright-Fisher SDE (1) with *ρ*(*t*) ≡ 1. At each time point between −0.4 and 0.0 in steps of 0:05, we simulated the sampling of 20 chromosomes.

We then ran the MCMC algorithm for 1, 000,000 generations, sampling every 1000 generations to obtain 1000 MCMC samples for each simulation. After discarding the first 500 samples from each MCMC run as burn-in, we computed the effective sample size of the allele age estimate using the R package coda [Plummer et al., 2006]. For the analysis of the simulations, we only included simulations that had an effective sample size greater than 150 for the allele age, resulting in retaining 744 out of 1000 simulations.

Because our MCMC analysis provides a full posterior distribution on parameter values, we summarized the results by computing the maximum *a posteriori* estimate of each parameter. We find that across the range of simulated *α*_1_ values, estimation is quite accurate (Figure 3A). There is some downward bias for large true values of *α*_1_, indicating the influence of the prior. On the other hand, the strength of selection in favor of the homozygote, *α*_2_, is less well estimated, with a more pronounced downward bias (Figure 3B). This is largely because most simulated alleles do not reach sufficiently high frequency for homozygotes to be common. Hence, there is very little information regarding the fitness of the homozygote. Allele age is estimated accurately, although there is a slight bias toward estimating a more recent age than the truth (Figure 3C).

**FIGURE 3.**
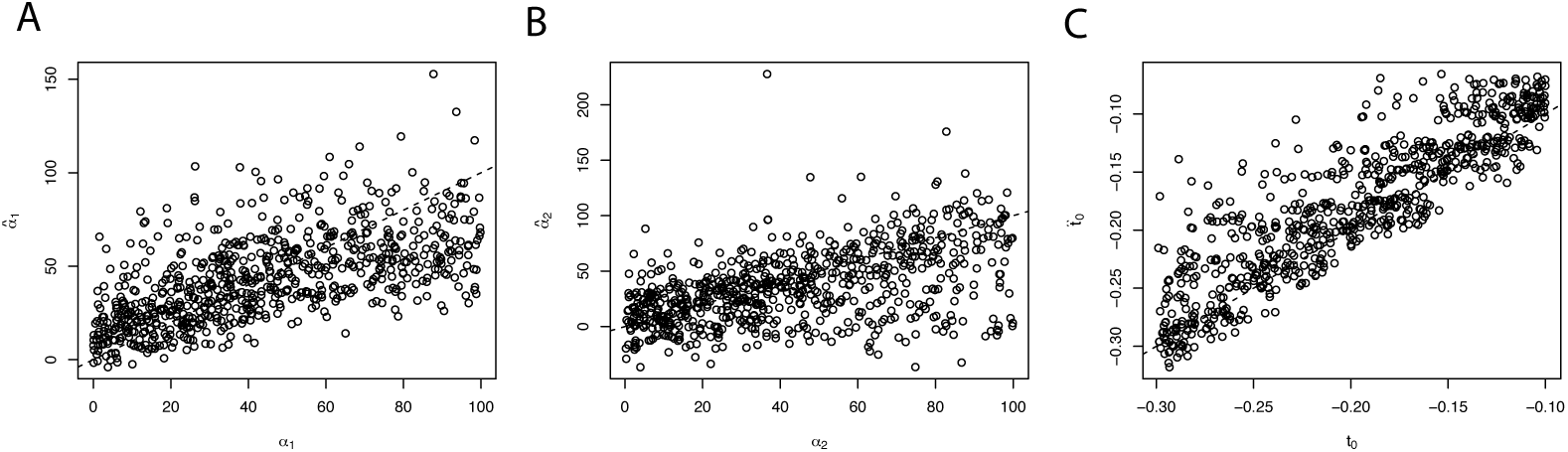
Maximum *a posterior* estimates of different parameters. Each panel shows the true value of a parameter on the *x*-axis, while the inferred value is on the *y*-axis. Dashed line is *y* = *x*.

When simulating under the horse demographic history, we drew 1000 allele ages with probability proportional to *ρ*(*t*) for *t* between 0.1 and 0.3 diffusion time units ago. Similarly to the simulations with constant demography, we drew *α*_1_ and *α*_2_ uniformly between 0 and 100), and then simulated allele frequency trajectories using an Euler approximation to (1) with p(t) given by the history inferred by Der Sarkissian et al. [2015]. The sampling scheme is identical to the constant demography simulations.

We ran our simulated data through two separate MCMC pipelines, one accounting for the true simulated demographic history, and the other assuming a constant population size. All other settings were identical to the analysis of the data simulated under constant demography. We retained MCMC runs where the sampling likelihood, path likelihood, a1 estimate, *α*_2_ estimate, and allele age estimate all had effective sample sizes greater than 50, resulting in 561 analyses retained from the inference with variable demography, 647 analyses retained from the inference with constant demography, and 454 analyses that were retained in both.

To quantify the overall impact of demographic model misspecification on parameter inference, we approximated the posterior root mean square error of a parameter (generically *θ*) by averaging over the posterior distribution,

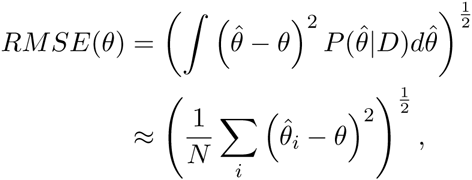

where the sum is over retained MCMC samples.

We found substantially smaller RMSE for inference of *α_1_* when demography is properly modeled (Figure 4). While inference of *α*_2_ was similar between the two models, there is somewhat larger RMSE when demography is incorrectly assumed to be constant. Interestingly, there seem to be two regimes of error in allele age estimation: for the most recent allele ages, modeling demography results in higher RMSE, while for more ancient ages, inferences with constant population size result in larger RMSE. These are likely caused by a particular feature of this demographic model, which is a very strong bottleneck inferred in the recent past. Because alleles are more likely to arise during periods of larger population size, accounting for demographic history extends the tail of the posterior distribution further into the past, when the population was larger.

**FIGURE 4.**
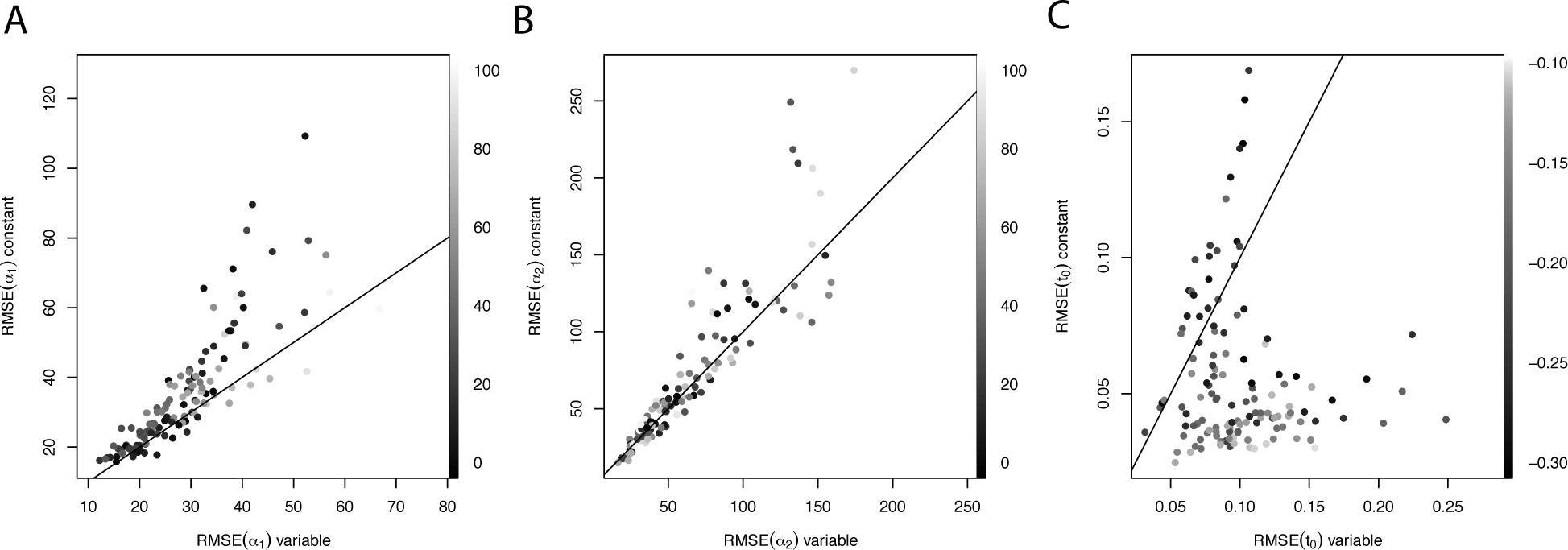
Comparison of root mean square error (RSME) when inference in performed with the proper (variable) demographic model on the *x*-axis compared to a misspecified constant demography model on the *y*-axis. Each point represents a single simulation, and points are colored according to simulated parameter value (scale on the right of each panel). Solid line is *y* = *x*.

### 3.2 Application to ancient DNA

We applied our approach to real data by reanalyzing the MC1R and ASIP data from Ludwig et al. [2009]. In contrast to earlier analyses of these data, we explicitly incorporated the demography of the domesticated horse, as inferred by Der Sarkissian et al. [2015], using a generation time of 8 years. Table 2 shows the sample configurations and sampling times corresponding to each locus, where diffusion units are scaled to 2*N*_0_, with *N*_0_ = 16000 being the most recent effective size reported by Der Sarkissian et al. [2015]. For comparison, we also analyzed the data assuming the population size has been constant at *N*_0_.

**TABLE 2.** Sample information for horse data. Diffusion time units are calculated assuming *N*_0_ = 2500 and a generation time of 5 years.

|  |  |  |  |  |  |  |
| --- | --- | --- | --- | --- | --- | --- |
| Sample time (years BCE) | 20,000 | 13,100 | 3,700 | 2,800 | 1,100 | 500 |
| Sample time (diffusion units) | 0.078 | 0.051 | 0.014 | 0.011 | 0.004 | 0.002 |
| Sample size | 10 | 22 | 20 | 20 | 36 | 38 |
| Count of ASIP alleles | 0 | 1 | 15 | 12 | 15 | 18 |
| Count of MC1R alleles | 0 | 0 | 1 | 6 | 13 | 24 |

With the MC1R locus, we found that posterior inferences about selection coefficients can be strongly influenced by whether or not demographic information is included in the analysis (Figure 5). Marginally, we see that incorporating demographic information results in an inference that *α*_1_ is larger than the constant-size model (MAP estimates of 267.6 and 74.1, with and without demography, respectively; Figure 5A), while *α*_2_ is inferred to be smaller (MAP estimates of 59.1 and 176.2, with and without demography, respectively; Figure 5B). This has very interesting implications for the mode of selection inferred on the MC1R locus. Recall that *α*_2_ > *α*_1_ > 0 is direction selection, in which the derived allele is always beneficial, *α*_2_ < *α*_1_ > 0 is overdominant selection, in which the heterozygote is favored, and *α*_2_ > *α*_1_ < 0 is underdominant selection, in which the heterozygote is disfavored. With constant demography, the trajectory of the allele is estimated to be shaped by positive directional selection (joint MAP, *α*_1_ = 87.6, *α*_2_ = 394.8; Figure 5C), while when demographic information is included, selection is inferred to act in an overdominant fashion (joint MAP, *α*_1_ = 262.5, *α*_2_ = 128.1; Figure 5D).

**FIGURE 5.**
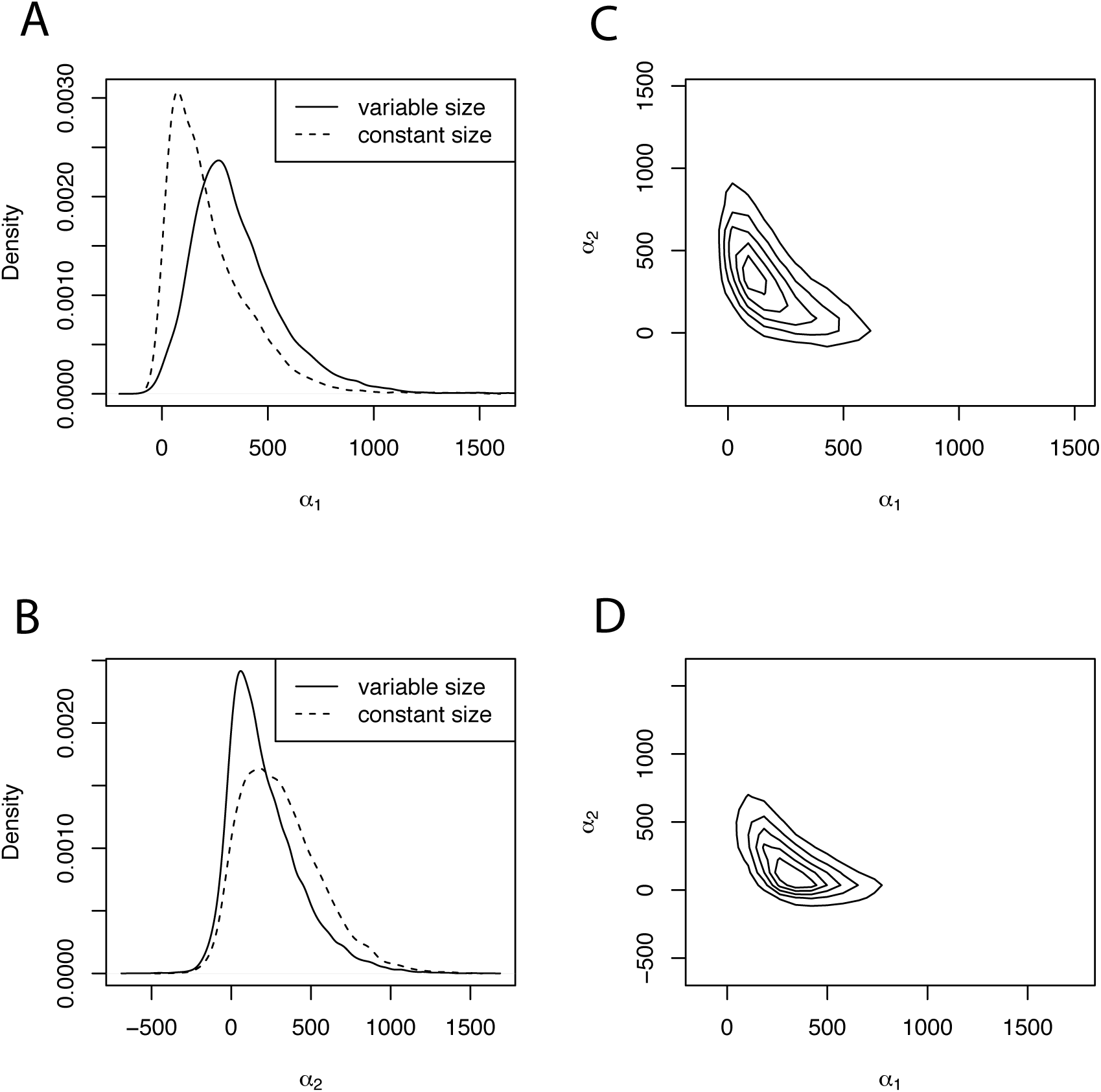
Posterior distributions of selection coefficients for the MC1R locus. Panels A and B show marginal distributions of *α*_1_ and *α*_2_, respectively, with the solid line indicating the posterior obtained from an analysis including the full demographic history, and the dotted line showing what would be inferred in a constant size population. Panels C and D show contour plots of the joint distribution of *α*_1_ and *α*_2_ without and with demography, respectively.

Incorporation of demographic history also has substantial impacts on the inferred distribution of allele ages (Figure 6). Most notably, the distribution of the allele age for MC1R is significantly truncated when demography is incorporated, in a way that correlates to the demographic events (Figure S1). While both the constant-size history and the more complicated history result in a posterior mode at approximately the same value of the allele age, the domestication bottleneck inferred by Der Sarkissian et al. [2015] makes it far less likely that the allele rose more anciently than the recent population expansion. Because the allele is inferred to be younger under the model incorporating demography, the strength of selection in favor of the homozygote must be higher to allow it to escape low frequency quickly and reach the observed allele frequencies. Hence, *α*_1_ is inferred to be much higher when demographic history is explicitly modeled.

**FIGURE 6.**
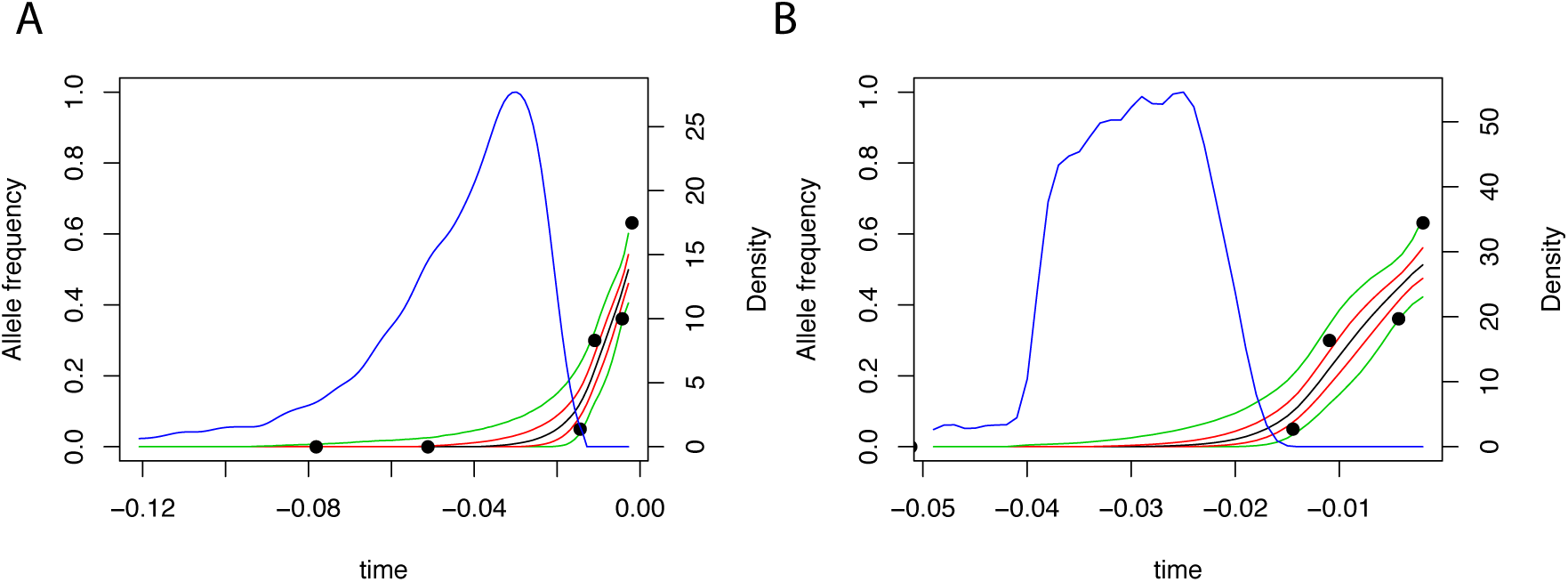
Posterior distribution on allele frequency paths for the MC1R locus. Each panel shows the sampled allele frequency data (filled circles), the point-wise median (black), 25 and 75% quantiles (red), and 5 and 95% quantiles (green) of the posterior distribution on paths, and the posterior distribution on allele age (blue). Panel A reports inference with constant demography, while panel B shows the result of inference with the full demographic history.

Incorporation of demographic history has an even more significant impact on inferences made about the ASIP locus (Figure 7). Most strikingly, while *α*_1_ is inferred to be very large without demography, it is inferred to be close to 0 when demography is incorporated (MAP estimates of 16.3 and 159.9 with and without demography, respectively; Figure 7A). On the other hand, inference of *α*_2_ is largely unaffected (MAP estimates of 34.7 and 39.8 with and without demography, respectively; Figure 7B). Interestingly, this has an opposite implication for the mode of selection compared to the results for the MC1R locus. With a constant-size demographic history, the allele is inferred to have evolved under overdominance (joint MAP, *α*_1_ = 153.3, *α*_2_ = 47; Figure 7C), but when the more complicated demography is modeled, the allele frequency trajectory is inferred to be shaped by positive, nearly additive, selection (joint MAP, *α*_1_ = 16.4, *α*_2_ = 46.8; Figure 7D).

**FIGURE 7.**
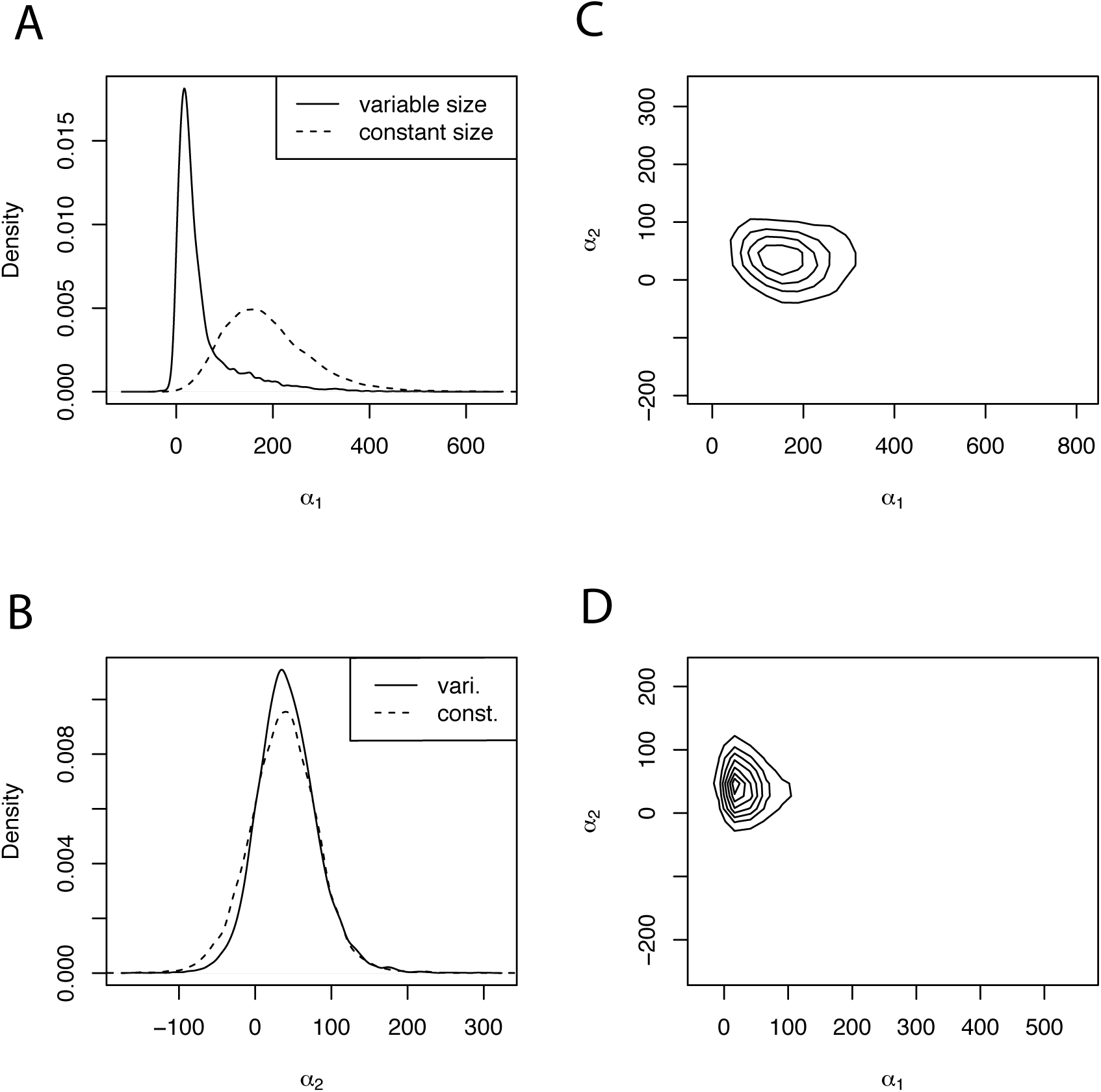
Posterior distributions of selection coefficients for the ASIP locus. Panels as in Figure 5.

Incorporating demography has a similarly opposite effect on inference of allele age (Figure 8). In particular, the allele is inferred to be much older when demography is modeled, and features a multi-modal posterior distribution on allele age, with each mode corresponding to a period of historically larger population size (Figure S2). Because the allele is inferred to be substantially older when demography is modeled, selection in favor of the heterozygote must have been weaker than would be inferred with the much younger age. Hence, the mode of selection switches from one of overdominance in a constant demography to one in which the homozygote is more fit than the heterozygote.

**FIGURE 8.**
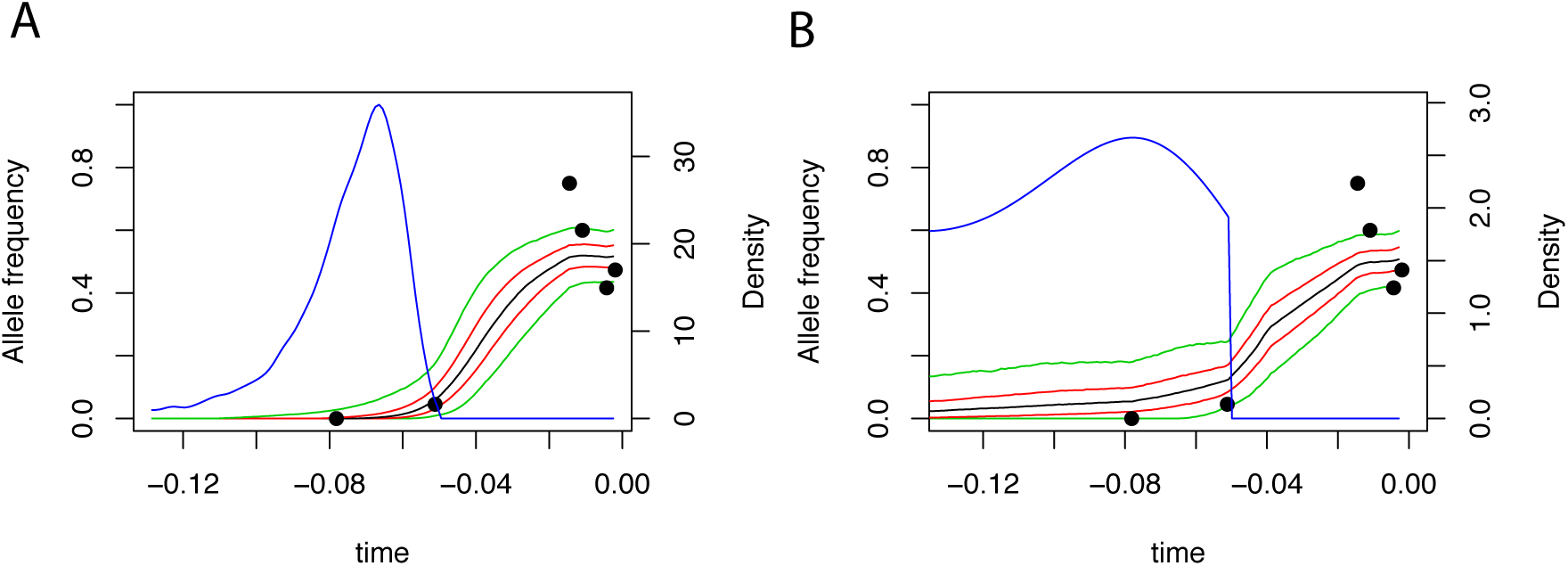
Posterior distribution on allele frequency paths for the ASIP locus. Panels are as in Figure 6.

## 4. DISCUSSION

Using DNA from ancient specimens, we have obtained a number of insights into evolutionary processes that were previously inaccessible. One of the most interesting aspects of ancient DNA is that it can provide a *temporal* component to evolution that has long been impossible to study. In particular, instead of making inferences about the allele frequencies, we can directly measure these quantities. To take advantage of this new data, we developed a novel Bayesian method for inferring the intensity and direction of natural selection from allele frequency time series. In order to circumvent the difficulties inherent in calculating the transition probabilities under the standard Wright-Fisher process of selection and drift, we used a data augmentation approach in which we learn the posterior distribution on allele frequency paths. Doing this not only allows us to efficiently calculate likelihoods, but provides an unprecedented glimpse at the historical allele frequency dynamics.

The key innovation of our method is to apply high-frequency path augmentation methods [Roberts and Stramer, 2001] to analyze genetic time series. The logic of the method is similar to the logic of a path integral, in which we average over all possible allele frequency trajectories that are consistent with the data [Schraiber, 2014]. By choosing a suitable reference probability distribution against which to compute likelihood ratios, we were able to adapt these methods to infer the age of alleles and properly account for variable population sizes through time. Moreover, because of the computational advantages of the path augmentation approach, we were able to infer a model of general diploid selection. To our knowledge, ours is the first work that can estimate both allele age and general diploid selection while accounting for demography.

Using simulations, we showed that our method performs well for strong selection and densely sampled time series. However, it is worth considering the work of Watterson [1979], who showed that even knowledge of the full trajectory results in very flat likelihood surfaces when selection is not strong. This is because for weak selection, the trajectory is extremely stochastic and it is difficult to disentangle the effects of drift and selection [Schraiber et al., 2013].

We also used simulations to test how misspecification of demographic history impacts inference. We saw substantially increased posterior root mean square error in inference of selection parameters if demographic history is misspecified. To examine the impact of demographic history in the context of real data, we then applied our method to a classic dataset from horses. We found that our inference of both the strength and mode of natural selection depended strongly on whether or not we incorporated demography. For the MC1R locus, a constant-size demographic model results in an inference of positive selection, while the more complicated demographic model inferred by Der Sarkissian et al. [2015] causes the inference to tilt toward overdominance, as well as a much younger allele age. In contrast, the ASIP locus is inferred to be overdominant under a constant-size demography, but the complicated demographic history results in an inference of positive selection, and a much older allele age.

These results stand in contrast to those of Steinrücken et al. [2014], who found that the most likely mode of evolution for both loci under a constant demographic history is one of overdominance. There are a several reasons for this discrepancy. First, we computed the diffusion time units differently, using *N*_0_ = 16000 and a generation time of 8 years, as inferred by Der Sarkissian et al. [2015], while Steinrücken et al. [2014] used *N*_0_ = 2500 (consistent with the bottleneck size found by Der Sarkissian et al. [2015]) and a generation time of 5 years. Hence, our constant-size model has far less genetic drift than the constant-size model assumed by Steinrücken et al. [2014]. This emphasizes the importance of inferring appropriate demographic scaling parameters, even when a constant population size is assumed. Secondly, we use MCMC to integrate over the distribution of allele ages, which can have a very long tail going into the past, while Steinruücken et al. [2014] assume a fixed allele age.

One key limitation of this method is that it assumes that the aDNA samples all come from the same, continuous population. If there is in fact a discontinuity in the populations from which alleles have been sampled, this could cause rapid allele frequency change and create spurious signals of natural selection. Several methods have been devised to test this hypothesis [Sjüdin et al., 2014], and one possibility would be to apply these methods to putatively neutral loci sampled from the same individuals, thus determining which samples form a continuous population. Alternatively, if our method is applied to a number of loci throughout the genome and an extremely large portion of the genome is determined to be evolving under selection, this could be evidence for model misspecification and suggest that the samples do not come from a continuous population.

An advantage of the method that we introduced is that it may be possible to extend it to incorporate information from linked neutral diversity. In general, computing the likelihood of neutral diversity linked to a selected site is difficult and many have used Monte Carlo simulation and importance sampling [Slatkin, 2001, Coop and Griffiths, 2004, Chen and Slatkin, 2013]. These approaches average over allele frequency trajectories in much same way as our method; however, each trajectory is drawn completely independently of the previous trajectories. Using a Markov chain Monte Carlo approach, as we do here, has the potential to ensure that only trajectories with a high posterior probability are explored and hence greatly increase the efficiency of such approaches.

## 5. ACKNOWLEDGMENTS

We are grateful for helpful comments and discussion with Yun Song, Matthias Steinrucken, and Anand Bhaskar during the conception and implementation of this work. We would also like thank 2 anonymous reviewers for their helpful comments that improved the clarity and thoroughness of this manuscript.

## 6. SOFTWARE AVAILABILITY

C++ software implementing the method described in this manuscript is freely available under a GNU Public License at https://github.com/Schraiber/selection.

## 7. APPENDIX

### 7.1 A proper posterior in the limit as the initial allele frequency approaches 0

For reasons that we explain in Subsection 2.4, we re-parametrize our model by replacing the path variable (*X_t_*)_*t*≥*t*_0__ with a deterministic time and space transformation of it (*Y_t_*)*_t_*>_0_ that takes values in the interval [0,π] with the boundary point 0 (resp. π) for (*Y_t_*)*_t_*≥_0_ corresponding to the boundary point 0 (resp. 1) for (*X_t_*)_*t*≥*t*_0__. The transformation producing (*Y_t_*)*_t_*≥_0_ is such that (*X_t_*)_*t*≥*t*_0__ can be recovered from (*Y_t_*)_*t*_≥_0_ and *t*_0_.

Implicit in our set-up is the initial frequency *x*_0_ at time *t*_0_ which corresponds to an initial value *y_0_* at time 0 of the transformed process (*Y_t_*)*_t_*≥_0_. For the moment, let us make the dependence on *y*_0_ explicit by including it in relevant notation as a superscript. For example, ℙ^y_0_^ (· | *α_1_, α_2_*,*t*_0_) is the prior distribution of (*Y_t_*)*_t_*≥_0_ given the specified values of the other parameters *α*_1_,*α*_2_,*t*_0_. We will construct a tractable “reference” process 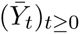 with distribution ℚ*^y^*^0^(·) such that the probability distribution ℙ^*y*_0_^(· | *α*_1_,*α*_2_,*t*_0_) has a density with respect to ℚ^*y*_0_^(·) – explicitly, ℚ^*y*0^(·) is the distribution of a Bessel(0) process started at location *y*_0_ at time 0. That is, there is a function Φ^*y*^_0_(·; *α*_1_,*α*_2_,*t*_0_) on path space such that

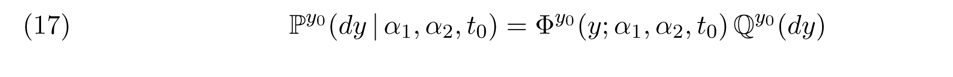

for a path (*y_t_*)_*t*≥0_. Assuming that π has a density with respect to Lebesgue measure which, with a slight abuse of notation, we also denote by π, the outcome of our Bayesian inferential procedure is determined by the ratios

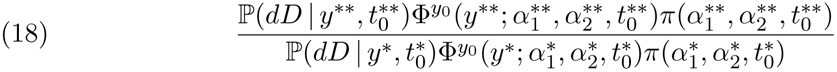

for pairs of augmented parameter values 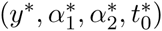 and 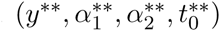 (*i.e*. the Metropolis-Hastings ratio).

Under the probability measure ℙ^*y*_0_^(· | *α*_1_,*α*_2_,*t*_0_), the process (*Y_t_*)_*t*≥0_ converges in distribution as *y*_0_ ↓ 0 (equivalently, *x*_0_ ↓ 0) to the trivial process that starts at location 0 at time 0 and stays there. However, for all *∊* > 0 the conditional distribution of (*Y_t_*)_*t*≥*∊*_ under the probability measure ℙ^*y*_0_^(· | *α*_1_,*α*_2_,*t*_0_) given the event {*Y_∊_* > 0} converges to a non-trivial probability measure as *y*_0_ ↓ 0. Similarly, the conditional distribution of the reference diffusion process 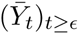 under the probability measure ℚ^*y*_0_^(·) given the event 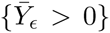 converges as *y*_0_ ↓ 0 to a non-trivial limit. There are *σ*-finite measures ℙ^0^(· | *α*_1_,*α*_2_,*t*_0_) and ℚ^0^(·) on path space that both have infinite total mass, are such that for any *∊* > 0 both of these measures assign finite, non-zero mass to the set of paths that are strictly positive at the time *∊*, and the corresponding conditional probability measures are the limits as *y*_0_ ↓ 0 of the conditional probability measures described above. Moreover, there is a function Φ^0^(·; *α*_1_,*α*_2_,*t*_0_) on path space such that

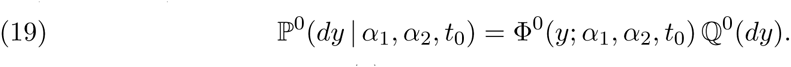

The posterior distribution (3) converges to

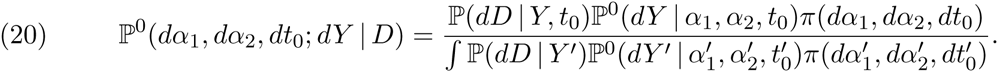

Thus, the limit as *y*_0_ ↓ 0 of a Bayesian inferential procedure for the augmented set of parameters can be viewed as a Bayesian inferential procedure with the improper prior ℙ^0^(*dY* | *α*_1_, *α*_2_, *t*_0_)*π*(*dα*_1_, *dα*_2_, *dt*_0_) for the parameters *Y*, *α*_1_,*α*_2_,*t*_0_. In particular, the limiting Bayesian inferential procedure is determined by the ratios

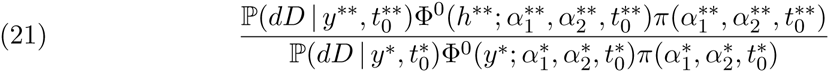

for pairs of augmented parameter values 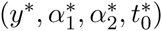 and (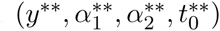.

### 7.2 The likelihood of the data and the path

Write *τ_i_* = *f* (*t_i_*). Note that *τ*_0_ = *f* (*t*_0_) = 0. Using equation (9), the density of the distribution of the transformed allele frequency process (*Y_t_*)_0≤*s*≤*τ_k_*_ against the reference distribution of the Bessel(0) process 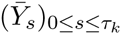 when 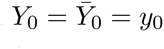 can be written

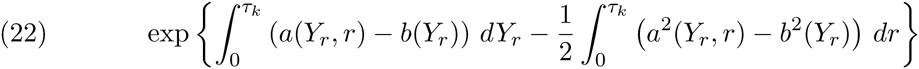

where

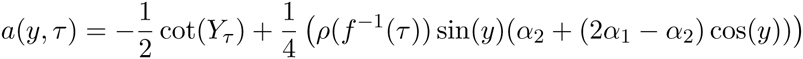

is the infinitesimal mean of the transformed Wright-Fisher process and

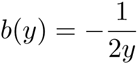

is the infinitesimal mean of the Bessel(0) process. However, as shown by Sermaidis et al. [2013], attempting to approximate the Itô integral in (22) using a discrete representation of the path can lead to biased estimates of the posterior distribution. Instead, consider the potential functions

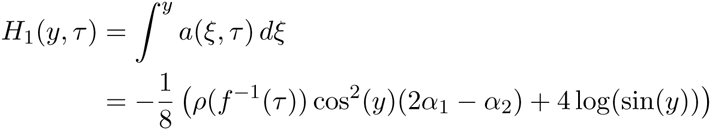

and

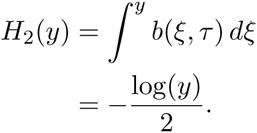

If we assume that *ρ* is continuous (not merely right continuous with left limits), then Itô’s lemma shows that we can write

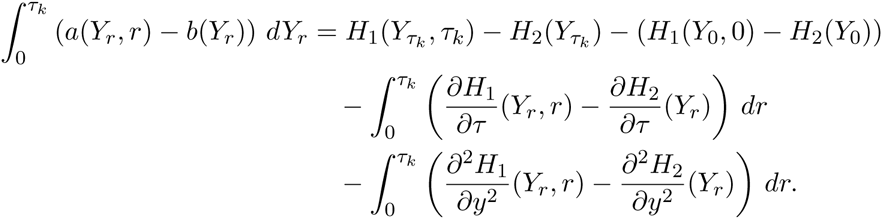

To generalize this to the case where *ρ* is right continuous with left limits, write

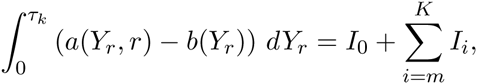

where *m* and *K* are defined in the main text,

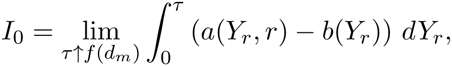

for *m* < *i* < *k*

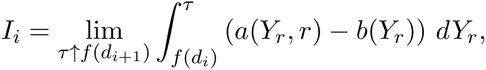

and

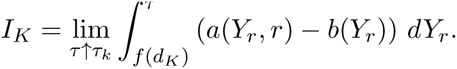

Itô’s lemma can then be applied to each segment in turn. Following the conversion of the Itô integrals into ordinary Lebesgue integrals, making the substitution *s* = *f*^−1^(*r*) results in the path likelihood displayed in (11).

### 7.3 Acceptance probability for an interior path update

When we propose a new path 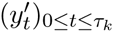 to update the current path (*y_t_*)_0≤*t*≤*τ_k_*_ which doesn’t hit the boundary, the new path agrees with the existing path outside some time interval [*υ*_1_,*υ*_2_], and has a new segment spliced in that goes from *y*_*υ*l_ at time *υ*_1_ to *y*_*υ*2_ at time *υ*_2_. The proposed new path segment comes from a Bessel(0) process over the time interval [*υ*_1_, *υ*_2_] that is pinned to take the values *y*_*υ*1_ and *y*_*υ*2_ at the end-points; that is, the proposed new piece of path is a bridge.

The ratio that determines the probability of accepting the proposed path is

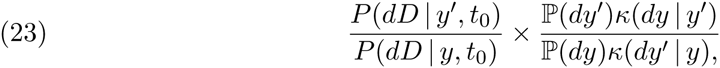

where *P*(· | *y*′,*t*_0_) and *P*(· | *y*,*t*_0_) give the probability of the observed allele counts given the transformed allele frequency paths and initial time *t*_0_, ℙ(·) is the distribution of the transformed Wright-Fisher diffusion starting from *y*_0_ > 0 at time 0 (that is, the distribution we have sometimes denoted by ℙ^*y*_0_^), the probability kernel κ(· | *y*) gives the distribution of the proposed path when the current path is *y*, and κ(· | *y*′) is similar. To be completely rigorous, the second term in the product in (23) should be interpreted as the Radon-Nikodym derivative of two probability measures on the product of path space with itself.

Consider a finite set of times 0 ≡ *τ*_0_ ξ *u*_0_ < *u*_1_ <… < *u*_ℓ_ ≡ *τ_k_*. Suppose that {*υ*_1_, *υ*_2_} ∈ {*u*_0_,…, *u*_ℓ_}, *υ*_1_ = *u_m_* and *υ*_2_ = *u_n_* for some *m* < *n*. Let (*y_t_*)_0≤*t*≤*τ_k_*_ and 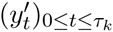 be two paths that coincide on [0, *υ*_1_] ∪ [*υ*_2_,*τ_k_*] = [*u*_0_,*u_m_*] ∪ [*u_n_*, *u_ℓ_*]. Write *P*(*x*,*y*; *s*,*t*) the transition density (with respect to Lebesgue measure) of the transformed Wright-Fisher diffusion from time *s* to time *t* and *Q*(*x*, *y*; *t*) for the transition density (with respect to Lebesgue measure) of the Bessel(0) process. Suppose that (*ξ*, *ζ*) is a pair of random paths with *P*((*ξ*,*ζ*) ∈ (*dy*,*dy*′)) = ℚ(*dy*)*κ*(*dy*′ | *y*). Then, writing 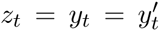 for *t* ∈ [0,*υ*_1_] ∪ [*υ*_2_, *τ_κ_*] = [*u*_0_,*u_m_*] ∪ [*u_n_*, *u_ℓ_*], we have

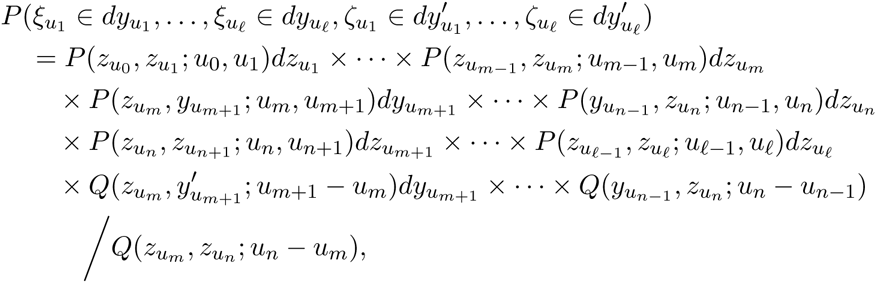

where the factor in the denominator arises because we are proposing *bridges* and hence conditioning on going from a fixed location at *υ*_1_ = *u_m_* to another fixed location at *υ*_2_ = *u_n_*.

Thus,

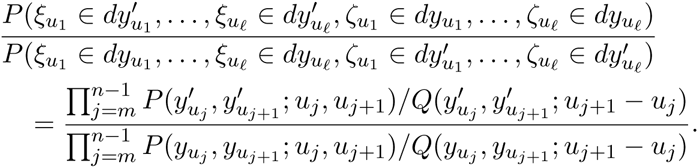

Therefore, the Radon-Nikodym derivative appearing in (23) is the ratio of Radon-Nikodym derivatives

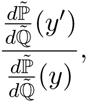

where 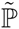 (resp. 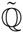) is the distribution of the transformed Wright-Fisher diffusion (resp. the Bessel(0) process) started at location 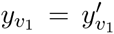 at time *υ*_1_ and run until time *υ*_2_. The formula (12) for the acceptance probability associated with an interior path update follows immediately.

The above argument was carried out under the assumption that the transformed initial allele frequency *y*_0_ was strictly positive and so all the measures involved were probability measures. However, taking *y*_0_ ↓ 0 we see that the formula (12) continues to hold. Alternatively, we could have worked directly with the measure ℙ^0^ in place of ℙ^*y*0^. The only difference is that we would have to replace *P*(*y*_0_, *y*; 0, *s*) by the density *ϕ*(*y*; 0, *s*) of an entrance law for ℙ^0^. That is, *ϕ*(*y*; 0, *s*) has the property that

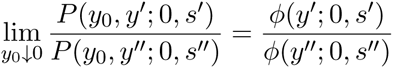

for all *y*′, *y*″ > 0 and *s*′, *s*″ > 0 so that

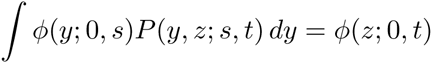

for 0 < *s* < *t*. Such a density, and hence the corresponding entrance law, is unique up to a multiplicative constant. In any case, it is clear that the choice of entrance law in the definition of ℙ^0^ does not affect the formula (12) as the entrance law densities “cancels out”.

### 7.4 Acceptance probability for an allele age update

The argument justifying the formula (13) for the probability of accepting a proposed update to the allele age *t*_0_ is similar to the one just given for interior path updates. Now, however, we have to consider replacing a path *y* that starts from *y*_0_ at time 0 and runs until time *f* (*t_k_*) with a path *y*′ that starts from *y*_0_ at time 0 and runs until time *f*′(*t_k_*). Instead of removing an internal segment of path and replacing it by one of the same length with the same values at the endpoints, we replace the initial segment of path that runs from time 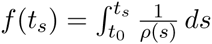 by one that runs from time 0 to time 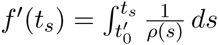 with 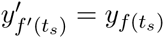.

By analogy with the previous subsection, we need to consider

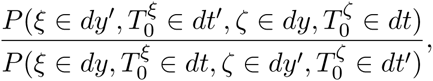

where *ξ* is a transformed Wright-Fisher process starting at *y*_0_ at time 0 and run to time 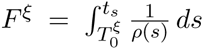, where 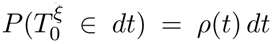, and conditional on *ξ*, *ζ* is a Bessel(0) bridge run from *y*_0_ at time 0 to *ξ_F^ξ^_* at time 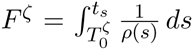, where 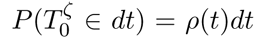 independent of *ξ* and 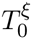.

Suppose that 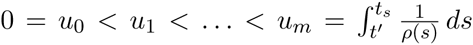 and 0 = *υ*_0_ < *υ*_1_ <… < *υ_n_* = 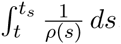. We have for 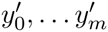 and *y*_0_,…,*y_n_* with 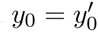 and 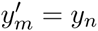 that

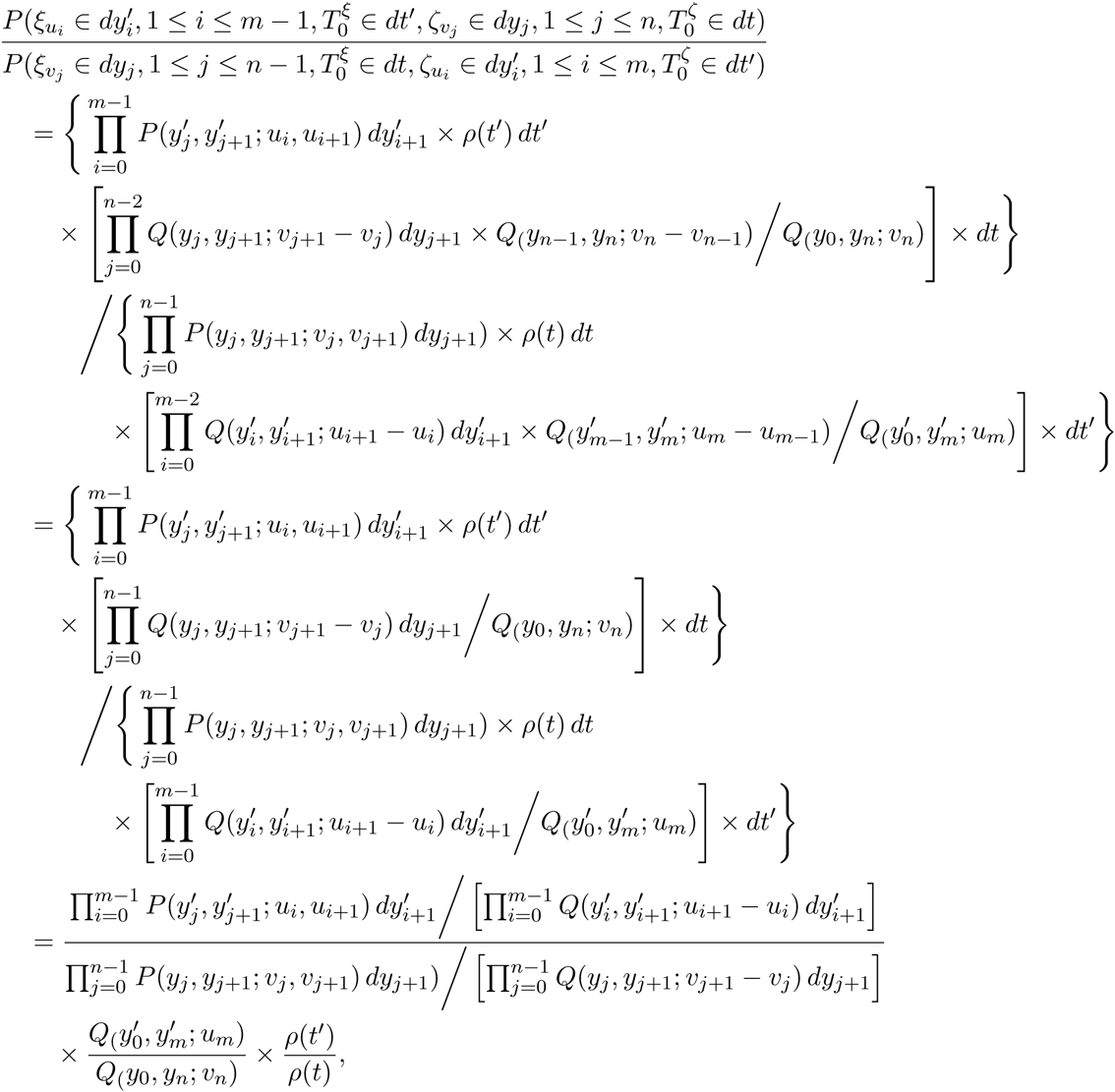

where the second equality follows from the fact that 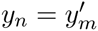.

Thus,

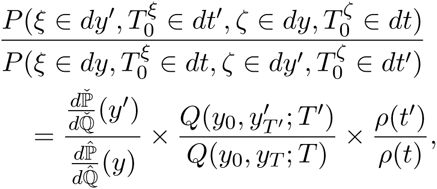

where 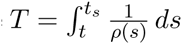 and 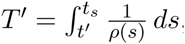 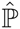(resp. 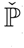) is the distribution of the transformed Wright-Fisher diffusion starting at location *y*_0_ at time 0 and run until time *T* (resp. *T*′), and 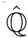 (resp. 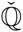) is the distribution of the Bessel(0) process starting at location *y*_0_ at time 0 and run until time *T* (resp. *T*′).

We have thusfar assumed that *y*_0_ is strictly positive. As in the previous subsection, we can let *y*_0_ ↓ 0 to get an expression in terms of Radon-Nikodym derivatives of *σ*-finite measures and the density *ψ*(*y*; *s*) of an entrance law for ℚ^0^. That is, *ψ*(*y*; *s*) has the property that

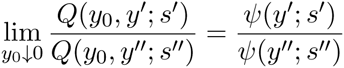

for all *y*′, *y*″ > 0 and *s*′, *s*″ > 0, so that

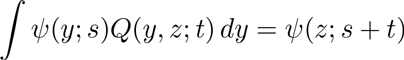

for *s*,*t* > 0. Up to an irrelevant multiplicative constant, *ψ* is given by the expression (14), and the formula (13) for the acceptance probability follows immediately.

### 7.5 Acceptance probability for a most recent allele frequency update

The derivation of formula (15) for the probability of accepting a proposed update to the most recent allele frequency is similar to those for the other acceptance probabilities (12) and (13), so we omit the details.

**FIGURE S1.**
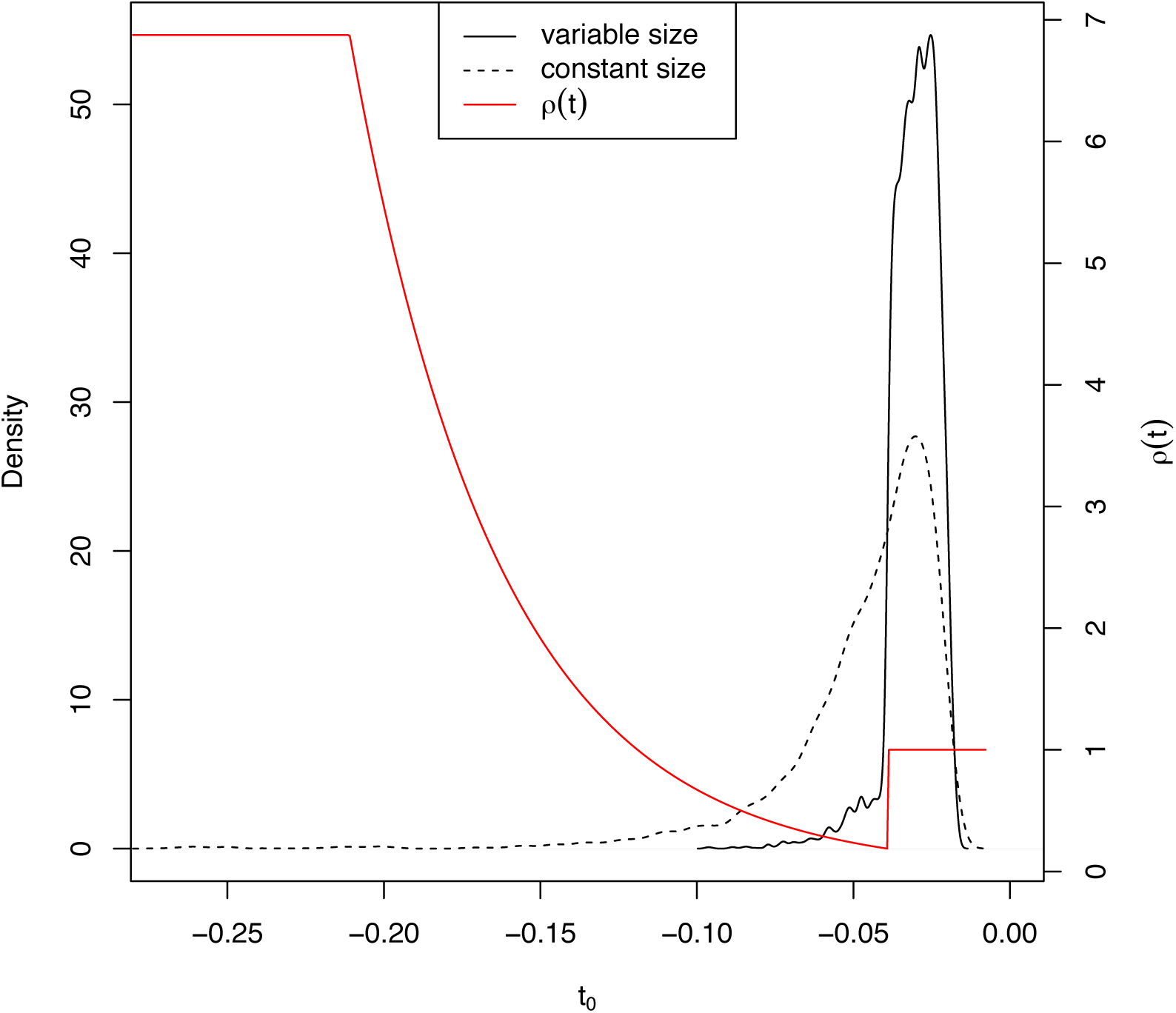
Influence of population size on age estimates of the MC1R locus. The solid and dashed lines show the posterior distribution on allele age with and without demography, respectively. In red, the demographic history inferred by Der Sarkissian et al. [2015].

**FIGURE S2.**
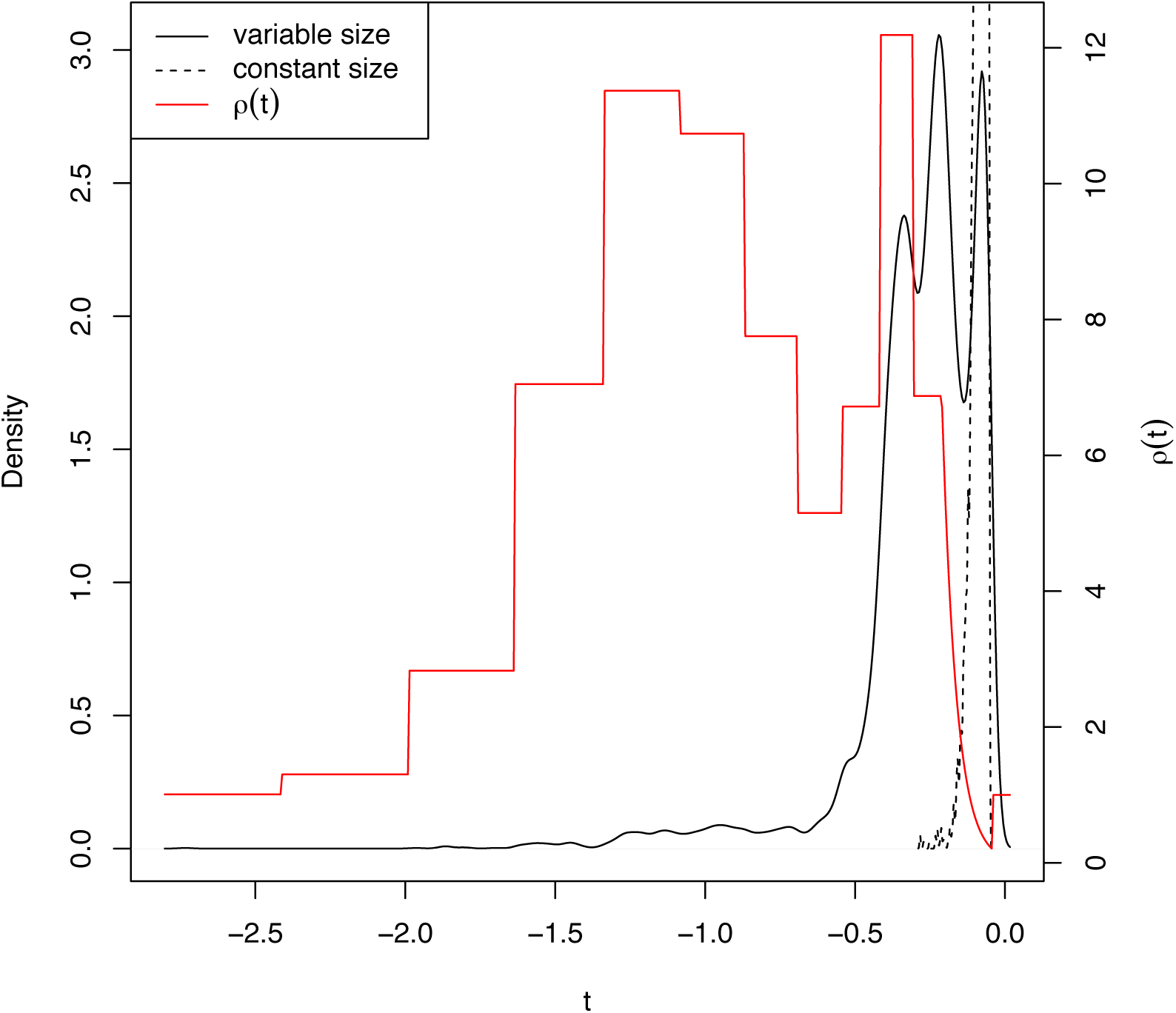
Influence of population size on age estimates of the ASIP locus. Data presented is as in Figure S1.

